# Broadly neutralizing antibodies overcome SARS-CoV-2 Omicron antigenic shift

**DOI:** 10.1101/2021.12.12.472269

**Authors:** Elisabetta Cameroni, Christian Saliba, John E. Bowen, Laura E. Rosen, Katja Culap, Dora Pinto, Laura A. VanBlargan, Anna De Marco, Samantha K. Zepeda, Julia di Iulio, Fabrizia Zatta, Hannah Kaiser, Julia Noack, Nisar Farhat, Nadine Czudnochowski, Colin Havenar-Daughton, Kaitlin R. Sprouse, Josh R. Dillen, Abigail E. Powell, Alex Chen, Cyrus Maher, Li Yin, David Sun, Leah Soriaga, Jessica Bassi, Chiara Silacci-Fregni, Claes Gustafsson, Nicholas M. Franko, Jenni Logue, Najeeha Talat Iqbal, Ignacio Mazzitelli, Jorge Geffner, Renata Grifantini, Helen Chu, Andrea Gori, Agostino Riva, Olivier Giannini, Alessandro Ceschi, Paolo Ferrari, Pietro Cippà, Alessandra Franzetti-Pellanda, Christian Garzoni, Peter J. Halfmann, Yoshihiro Kawaoka, Christy Hebner, Lisa A. Purcell, Luca Piccoli, Matteo Samuele Pizzuto, Alexandra C. Walls, Michael S. Diamond, Amalio Telenti, Herbert W. Virgin, Antonio Lanzavecchia, David Veesler, Gyorgy Snell, Davide Corti

## Abstract

The recently emerged SARS-CoV-2 Omicron variant harbors 37 amino acid substitutions in the spike (S) protein, 15 of which are in the receptor-binding domain (RBD), thereby raising concerns about the effectiveness of available vaccines and antibody therapeutics. Here, we show that the Omicron RBD binds to human ACE2 with enhanced affinity relative to the Wuhan-Hu-1 RBD and acquires binding to mouse ACE2. Severe reductions of plasma neutralizing activity were observed against Omicron compared to the ancestral pseudovirus for vaccinated and convalescent individuals. Most (26 out of 29) receptor-binding motif (RBM)-directed monoclonal antibodies (mAbs) lost in vitro neutralizing activity against Omicron, with only three mAbs, including the ACE2-mimicking S2K146 mAb^1^, retaining unaltered potency. Furthermore, a fraction of broadly neutralizing sarbecovirus mAbs recognizing antigenic sites outside the RBM, including sotrovimab^2^, S2X259^3^ and S2H97^4^, neutralized Omicron. The magnitude of Omicron-mediated immune evasion and the acquisition of binding to mouse ACE2 mark a major SARS-CoV-2 mutational shift. Broadly neutralizing sarbecovirus mAbs recognizing epitopes conserved among SARS-CoV-2 variants and other sarbecoviruses may prove key to controlling the ongoing pandemic and future zoonotic spillovers.

## INTRODUCTION

The evolution of RNA viruses can result in immune escape and modulation of binding to host receptors^5^. Previous SARS-CoV-2 variants of concern (VOC) have developed resistance to neutralizing antibodies, including some clinical antibodies used as therapeutics^6-9^. The B.1.351 (Beta) VOC demonstrated the greatest magnitude of immune evasion from serum neutralizing antibodies^6,7^, whereas B.1.617.2 (Delta) quickly outcompeted all other circulating isolates through acquisition of mutations that enhanced transmission and pathogenicity^10-13^ and eroded neutralizing antibody responses^10^.

The Omicron (B.1.1.529.1) variant was first detected in November 2021, whereupon it was immediately declared by the WHO as a VOC and quickly rose in frequency worldwide (**Extended Data Fig. 1**). Strikingly, analysis of the substitutions within the Omicron variant showed substantial changes from any previously described SARS-CoV-2 isolates, including 37 S protein mutations in the predominant haplotype (**Fig. 1a-b** and **Extended Data Fig. 1-4**). Fifteen of the Omicron mutations are clustered in the RBD, which is the major target of neutralizing antibodies upon infection and vaccination^14,15^, suggesting that Omicron may escape infection- and vaccine-elicited Abs and therapeutic mAbs. Nine of these mutations map to the receptor-binding motif (RBM) which is the RBD subdomain directly interacting with the host receptor, ACE2^16^.

**Fig. 1.**
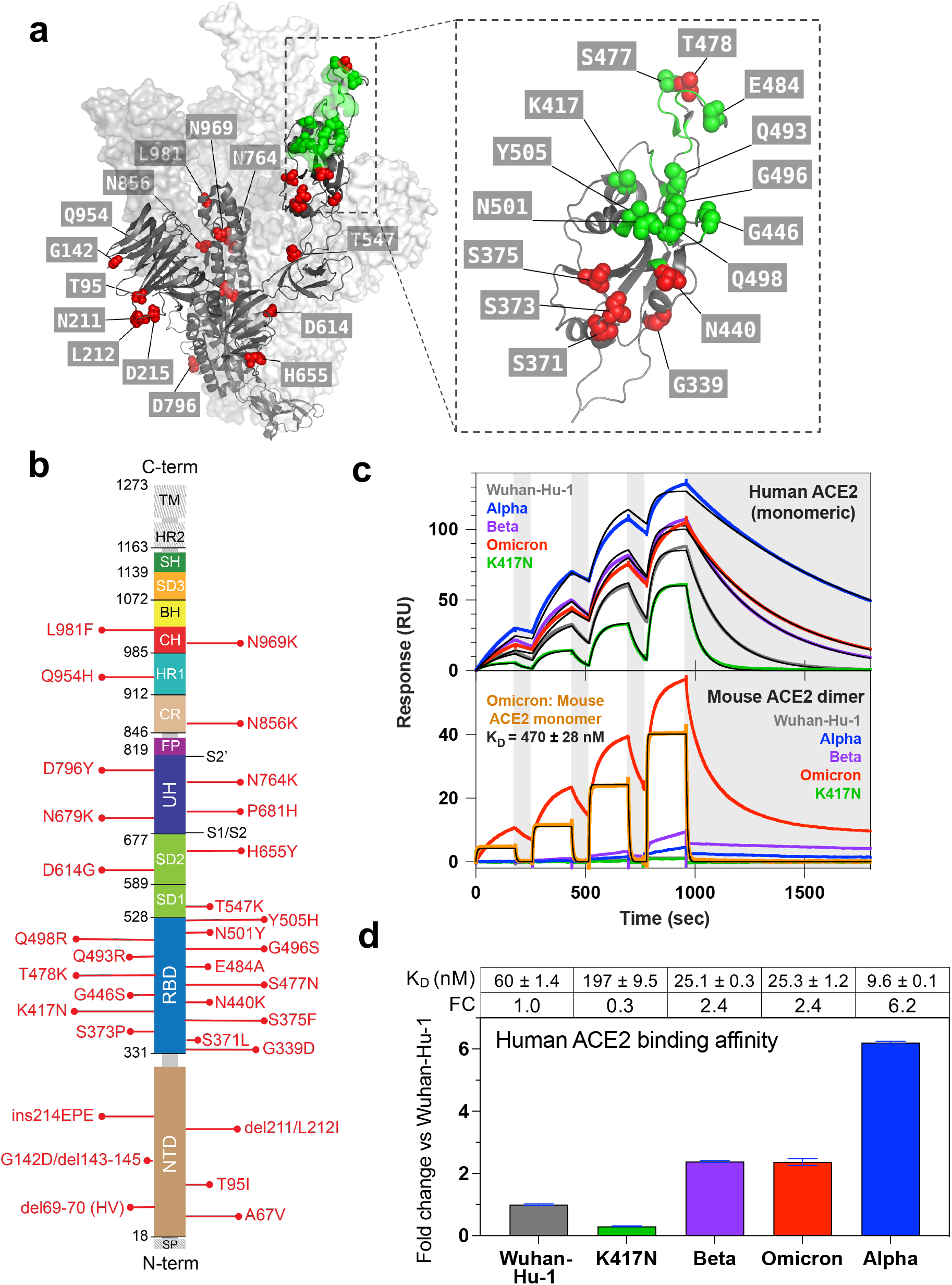
Omicron RBD shows increased binding to human ACE2 and gains binding to murine ACE2. **a**, SARS-CoV-2 S in fully open conformation (PDB: 7K4N) with positions of mutated residues in Omicron highlighted on one protomer in green or red spheres in or outside the ACE2 footprint (ACE2), respectively. RBM is defined by a 6 Å cutoff in the RBD-ACE2 interface^42^. **b**, Omicron mutations are shown in a primary structure of SARS-CoV-2 S with domains and cleavage sites highlighted. **c**, Single-cycle kinetics SPR analysis of ACE2 binding to five RBD variants. ACE2 is injected successively at 11, 33, 100, and 300 nM (human) or 33, 100, 300, and 900 nM (mouse); monomeric and dimeric mouse ACE2 were tested. Black curves show fits to a 1:1 binding model. White and gray stripes indicate association and dissociation phases, respectively. **d**, Quantification of human ACE2 binding data. Reporting average ± standard deviation of three replicates.

Preliminary reports indicated that the neutralizing activity of plasma from Pfizer-BioNTech BNT162b2 vaccinated individuals is severely reduced against SARS-CoV-2 Omicron^17,18^, documenting a substantial, albeit not complete, escape from mRNA vaccine-elicited neutralizing antibodies. Another report also showed that vaccine effectiveness against symptomatic disease with the Omicron variant is significantly lower than with the Delta variant^19^. The potential for booster doses to ameliorate this decline in neutralization is still being explored. In addition, the neutralizing activity of several therapeutic mAbs was shown to be decreased or abolished against SARS-CoV-2 Omicron^18,20^.

To understand the consequences of the unprecedented number of mutations found in Omicron S, we employed a pseudovirus assay to study neutralization mediated by monoclonal and polyclonal antibodies as well as surface plasmon resonance to measure binding of RBD to human and animal ACE2 receptors.

## RESULTS

### Omicron RBD binds with increased affinity to human ACE2 and gains binding to mouse ACE2

The unprecedented number of substitutions found in the Omicron genome raises questions about its origin. Twenty-three out of the 37 Omicron S amino acid mutations have been individually observed previously in SARS-CoV-2 variants of interest (VOI), VOC, or other sarbecoviruses, whereas the remaining 14 substitutions have not been described before in any SARS-CoV-2 isolates (**Extended Data Fig. 5a**). Analysis of the GISAID database^21^ indicates that there were rarely more than 10-15 Omicron S mutations present in a given non-Omicron haplotype or Pango lineage (**Extended Data Fig. 5b, c and d)**. While we have not formally assessed the possibility of recombination events, persistent replication in immunocompromised individuals or inter-species ping-pong transmission^5^ are possible scenarios for the rapid accumulation of mutations that could have been selected based on fitness and immune evasion.

To assess the latter scenario, we investigated whether RBD mutations found in Omicron may have resulted from adaptation of SARS-CoV-2 to animal receptors. To this end, we tested RBD binding to mouse, American mink, and pangolin ACE2 receptors by surface plasmon resonance (SPR) (**Fig. 1c** and **Extended Data Fig. 6**). Omicron bound mouse, but not mink or pangolin, ACE2 whereas the Wuhan-Hu-1, Beta, Alpha and K417N RBDs did not recognize any of these three ACE2 orthologues in our assay. Acquisition of mouse ACE2 binding is likely explained by the Q493R substitution which is very similar to the Q493K mutation isolated upon mouse adaptation of SARS-CoV-2^22^.

Several of the Omicron RBD mutations are found at positions that are key contact sites with human ACE2, such as K417N, Q493K and G496S^23,24^. Except for N501Y, which increases ACE2 binding affinity by 6-fold^25^, all other substitutions were shown by deep mutational scanning (DMS) to reduce binding to human ACE2 individually^26^, resulting in a marked predicted decrease of affinity (**Extended Data Table 1**). However, we found that the Omicron RBD has a 2.4-fold increased binding affinity to human ACE2 (**Fig. 1d**), suggesting epistasis of the full constellation of RBD mutations.

Collectively, these findings suggest that mutations in the RBD of Omicron may have enabled adaptation to rodents as well as contributed to potentially increased transmission in humans.

### Omicron escapes polyclonal plasma neutralizing antibodies

To investigate the magnitude of immune evasion mediated by the 37 mutations present in Omicron S, we determined plasma neutralizing activity against Wuhan-Hu-1 S and Omicron S VSV pseudoviruses in different cohorts of convalescent patients or individuals vaccinated with six major COVID-19 vaccines (mRNA-1273, BNT162b2, AZD1222, Ad26.COV2.S, Sputnik V and BBIBP-CorV) (**Fig. 2, Extended Data Figure 7-8** and **Extended Data Table 2**).

**Fig. 2.**
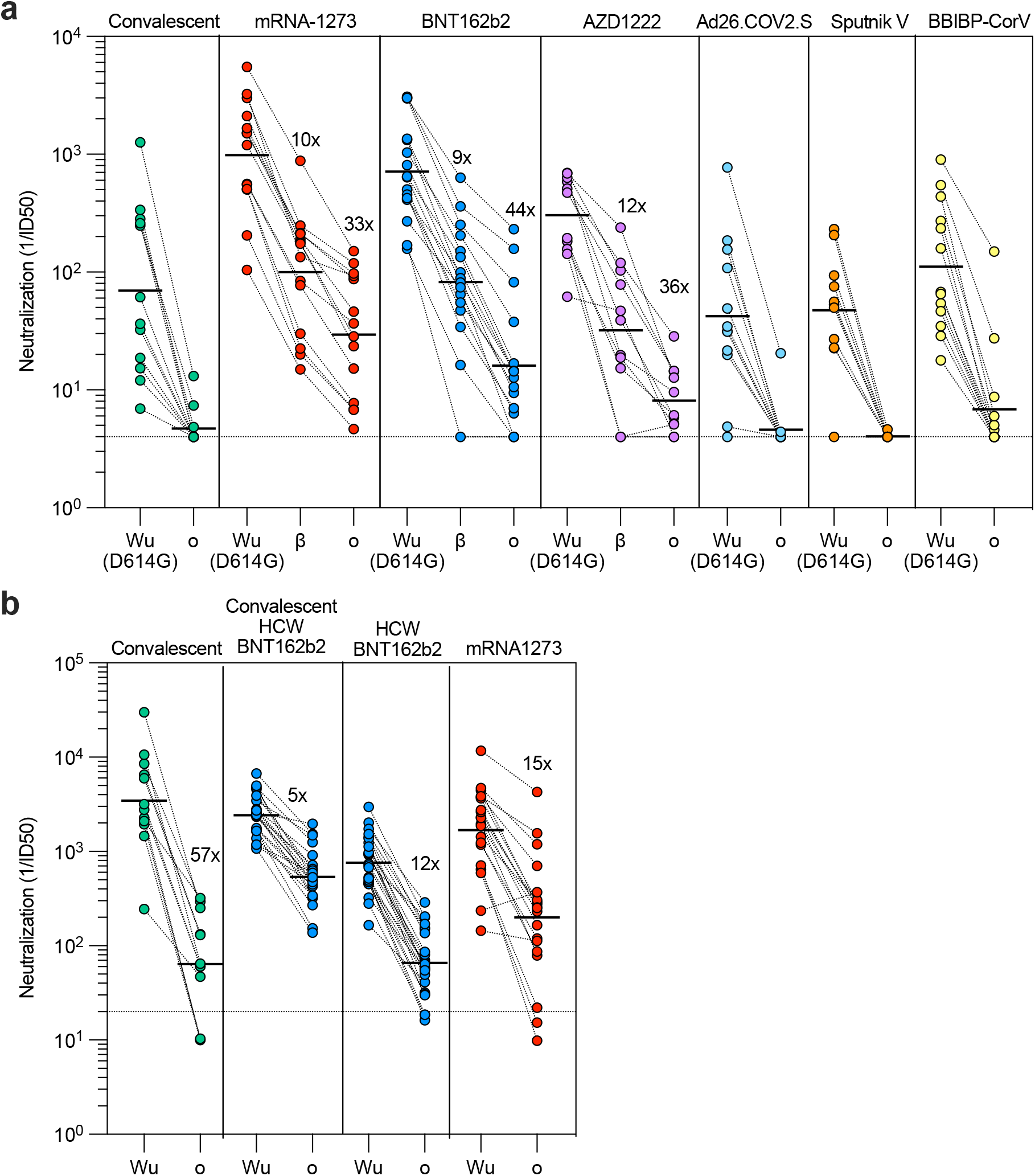
Neutralization of Omicron SARS-CoV-2 VSV pseudovirus by plasma from COVID-19 convalescent and vaccinated individuals. Plasma neutralizing activity in COVID-19 convalescent or vaccinated individuals (mRNA-1273, BNT162b2, AZD1222, Ad26.COV2.S (single dose), Sputnik V and BBIBP-CorV). **a**, Pairwise neutralizing antibody titers (ID50) against Wuhan-Hu-1 (D614G), Beta and Omicron VOC. Vero E6-TMPRSS2 used as target cells. Shown one representative experiment out of 2. **b**, Pairwise neutralizing antibody titers of plasma (ID50) against Wuhan-Hu-1 and Omicron VOC. 11 out of 12 convalescent donors were hospitalized for COVID-19. Vero E6 used as target cells. Data are average of n = 2 replicates. Line, geometric mean of 1/ID50 titers. HCW, healthcare workers; Wu, Wuhan-Hu-1; o, Omicron VOC, b, Beta VOC. Enrolled donors’ demographics provided in **Extended Data Table 2**.

Convalescent patients and individuals vaccinated with Ad26.COV2.S (single dose), Sputnik V or BBIBP-CorV had no neutralizing activity against Omicron except for one Ad26.COV2.S and three BBIBP-CorV vaccinees (**Fig. 2a-b**). Individuals vaccinated with mRNA-1273, BNT162b2, and AZD1222 displayed higher neutralization against Wuhan-Hu-1 and retained activity against Omicron with a decrease of 33-, 44- and 36-fold, respectively (**Fig. 2a**). Interestingly, this decrease was less pronounced for vaccinated individuals who were previously infected (5-fold) (**Fig. 2b**) consistent with broadening of antibody responses as a consequence of affinity maturation driven by multiple antigenic stimulations^27-29^. Collectively, these findings demonstrate a substantial and unprecedented reduction in plasma neutralizing activity against Omicron versus the ancestral virus, that in several cases may fall below protective titers^30^.

### Broadly neutralizing sarbecovirus antibodies retain activity against SARS-CoV-2 Omicron

Neutralizing mAbs with demonstrated in vivo efficacy in prevention or treatment of SARS-CoV-2^31-41^ can be divided into two groups based on their ability to block S binding to ACE2. Out of the eight currently authorized or approved mAbs, seven (LY-CoV555, LY-CoV016, REGN10933, REGN10933, COV2-2130, COV2-2196 and CT-P59; all synthesized based on publicly available sequences, respectively) block binding of S to ACE2 and are often used in combination^9^. These mAbs bind to epitopes overlapping with the RBM (**Fig. 3a**) which is structurally and evolutionary plastic^42^, as illustrated by the accumulation of mutations throughout the pandemic and the diversity of this subdomain among ACE2-utilizing sarbecoviruses^43^. Combining two such ACE2 blocking mAbs provides greater resistance to variant viruses that carry RBM mutations^32^. The second class of mAbs, represented by sotrovimab, do not block ACE2 binding but neutralize SARS-CoV-2 by targeting non-RBM epitopes shared across many sarbecoviruses, including SARS-CoV^4,44^.

**Fig. 3.**
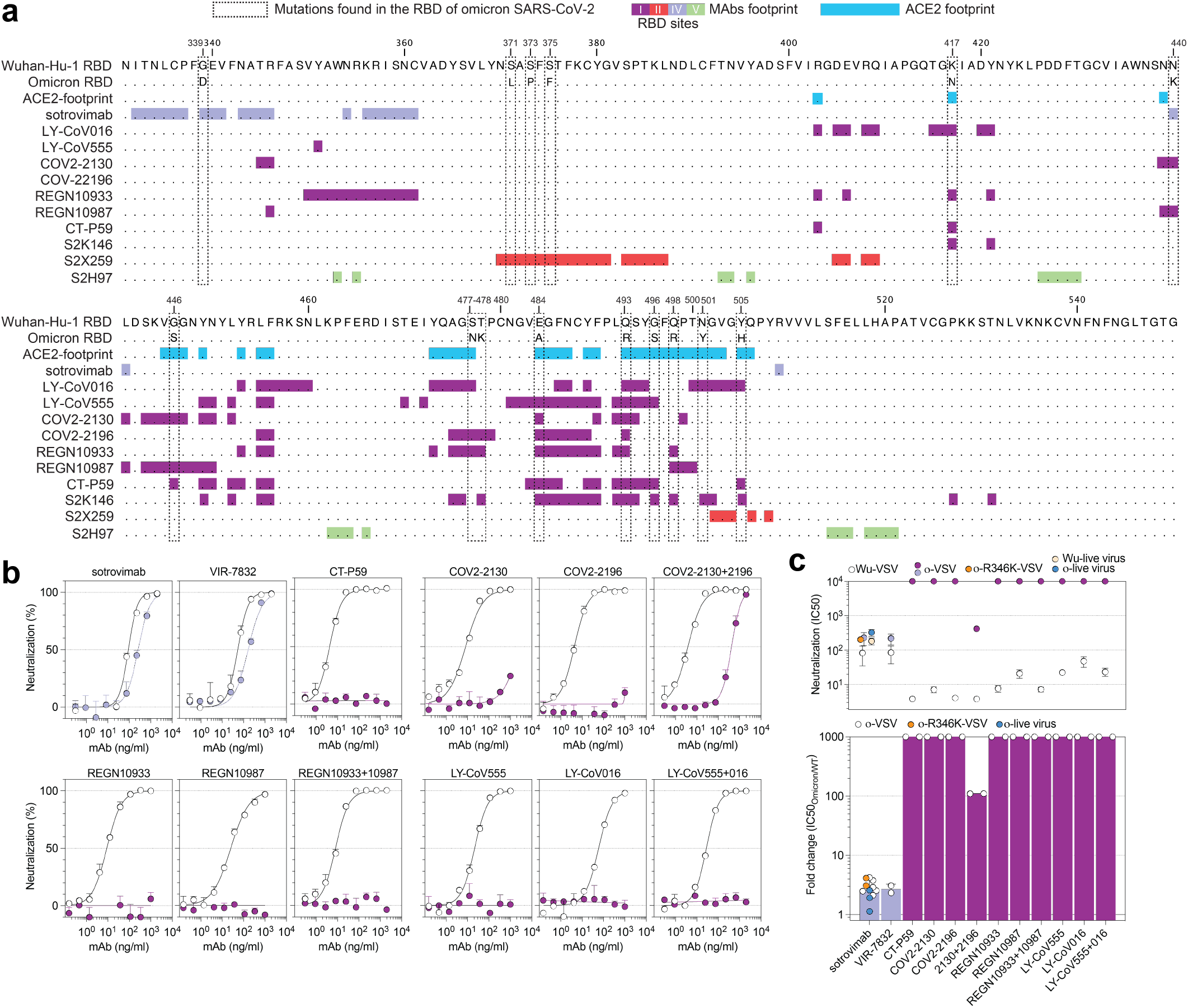
Neutralization of Omicron SARS-CoV-2 VSV pseudovirus by clinical-stage mAbs. **a**, RBD sequence of SARS-CoV-2 Wuhan-Hu-1 with highlighted footprints of ACE2 (light blue) and mAbs (colored according to the RBD antigenic site recognized). Omicron RBD is also shown, and amino acid substitutions are boxed. **b**, Neutralization of SARS-CoV-2 VSV pseudoviruses carrying Wuhan-Hu-1 (white) or Omicron (colored as in **Fig. 4b**) S proteins by clinical-stage mAbs. Data are representative of one independent experiment out of two. Shown is the mean ± s.d. of 2 technical replicates. **c**, Mean IC50 values for Omicron (colored as in Fig. 4b) and Wuhan-Hu-1 (white) (top panel), and mean fold change (bottom panel). Vero E6 used as target cells. Shown in yellow and blue is also neutralization of live virus by sotrovimab (WA1/2020 and hCoV-19/USA/WI-WSLH-221686/2021 isolates, respectively). Non-neutralizing IC50 titers and fold change were set to 10^4^ and 10^3^, respectively. Orange dots for sotrovimab indicate neutralization of Omicron carrying R346K. Data are representative of n = 2 to 6 independent experiments.

Here, we compared the in vitro neutralizing activity of therapeutic mAbs from these two groups against Wuhan-Hu-1 S and Omicron S using VSV pseudoviruses. Although sotrovimab had 3-fold reduced potency against Omicron (a similar potency was also measured against Omicron-R346K VSV pseudoviruses), all other (RBM-specific) mAbs completely lost their neutralizing activity with the exception of the cocktail of COV2-2130 and COV2-2196 for which we determined a ∼200-fold reduced potency (**Fig. 3b-c**). These findings are consistent with two recent reports^18,20^ and, together with serological data, support the notion of Omicron antigenic shift. Of note, sotrovimab also showed a less than 2-fold reduction in neutralizing activity against live Omicron SARS-CoV-2 as compared to the WAI/2020 D614G isolate (**Fig. 3c** and **Extended Data Fig. 9**), consistently with a recent report on S309, parent mAb of sotrovimab^45^.

We next tested a larger panel of 36 neutralizing NTD- or RBD-specific mAbs for which the epitope has been characterized structurally or assigned to a given antigenic site through competition studies^3,4,10,14,46,47^ (**Fig. 4a, Extended Data Table 2** and **Extended Data Fig. 10**). The four NTD-specific antibodies completely lost activity against Omicron, in line with the presence of several mutations and deletions in the NTD antigenic supersite^8,25^. Three out of the 22 mAbs targeting the RBD antigenic site I (RBM) retained potent neutralizing activity against Omicron, including S2K146, which binds the RBD of SARS-CoV-2, SARS-CoV and other sarbecoviruses through ACE2 molecular mimicry^1^. Out of the nine mAbs specific for the conserved RBD site II^4^ (class 4 mAbs), only S2X259^3^ retained activity against Omicron, whereas neutralization was decreased by more than 10-fold or abolished for the remaining mAbs. Finally, neutralization of Omicron was also retained with the S2H97 mAb, which recognizes the highly conserved cryptic site V. The panel of 44 mAbs tested in this study represent members of each of the four classes of broadly neutralizing sarbecovirus mAbs, defined by their cognate RBD binding sites (site I, II, IV and V). Our findings show that member(s) of each of the four classes can retain Omicron neutralization: S2K146, S2X324 and S2N28 targeting site I, S2X259 targeting site II, sotrovimab targeting site IV, and S2H97 targeting site V (**Fig. 4b**). Several of these mAbs cross-react with and neutralize sarbecoviruses beyond the SARS-CoV-2 clade 1b^1,3,4^, confirming the notion that targeting conserved epitopes can result not only in breadth but also in protection against viral evolution.

**Fig. 4.**
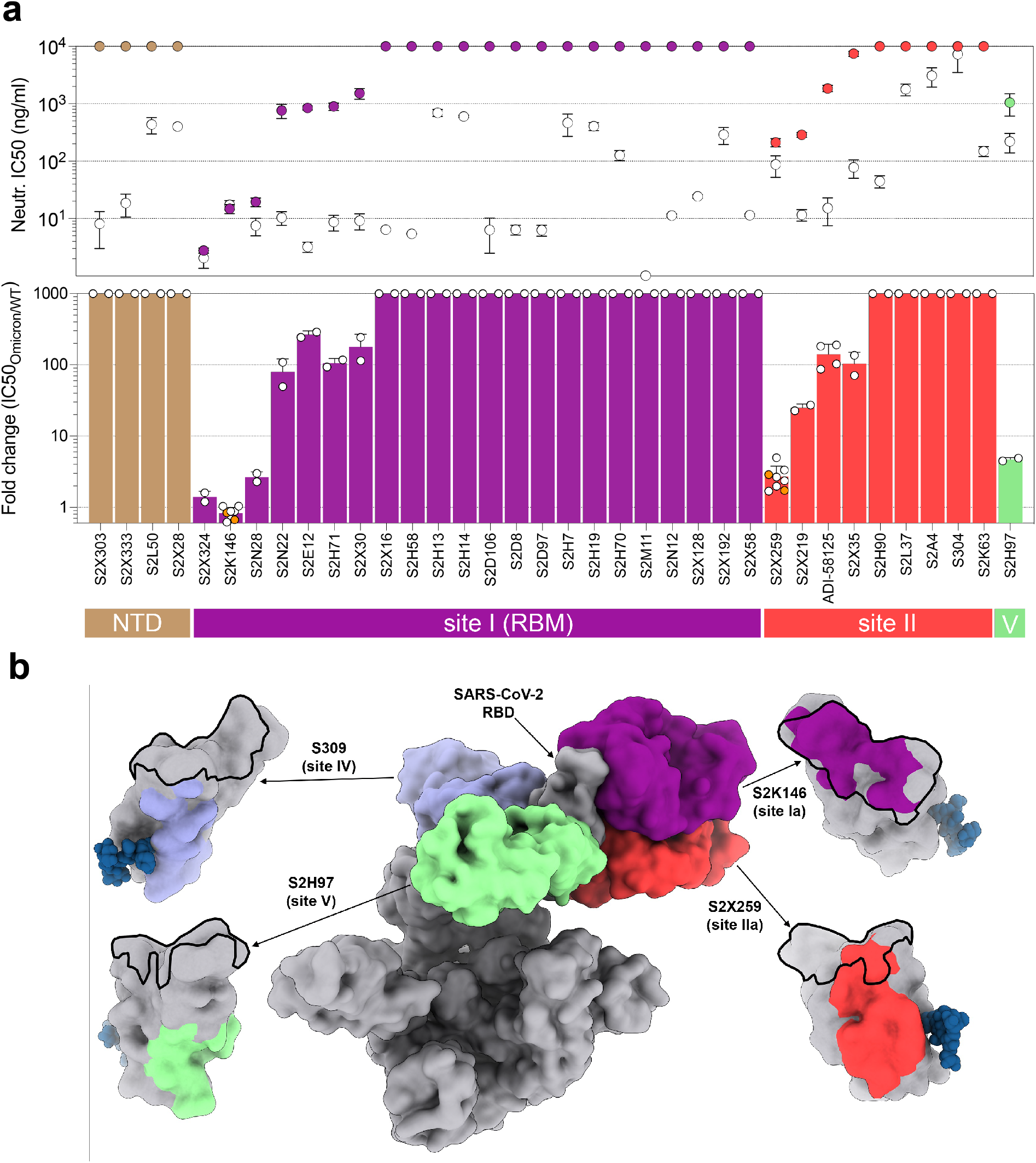
Neutralization of Omicron SARS-CoV-2 VSV pseudovirus by monoclonal antibodies. **a**, Mean IC50 values for Omicron (colored as in b) and Wuhan-Hu-1 (white) (top panel), and mean fold change (bottom panel) for 4 NTD mAbs and 32 RBD mAbs. Non-neutralizing IC50 titers and fold change were set to 10^4^ and 10^3^, respectively. Orange dots for S2K146 and S2X259 indicate neutralization of Omicron carrying R346K. Vero E6 used as target cells. Data are representative of n = 2 to 6 independent experiments. **b**, The RBD sites targeted by 4 mAbs cross-neutralizing Omicron are annotated and representative antibodies (the Fv region) bound to S are shown as a composite. Colored surfaces on the RBD depict the epitopes and the RBM is shown as a black outline.

## Discussion

The staggering number of substitutions present in Omicron S marks a dramatic shift in antigenicity and is associated with immune evasion of unprecedented magnitude for SARS-CoV-2 and a putative broadening of tropism. While influenza antigenic shift is defined as genetic reassortment of the RNA genome segments, the mechanism of accumulation of a large number of mutations in SARS-CoV-2 Omicron S remains to be established. Although recombination events are a coronavirus hallmark^48^, we and others^49^ propose that the Omicron shift may result from extensive viral replication in immunodeficient hosts^50-52^ or from inter-species ping-pong transmission^5^ between humans and rodents, as previously described for minks^53,54^.

Consistent with the variable decrease in plasma neutralizing antibody titers, we found that only 6 out of a panel of 44 neutralizing mAbs retained potent neutralizing activity against Omicron. These mAbs recognize four antigenic sites in the RBD that are conserved in Omicron and other sarbecoviruses. Strikingly, we found three potent neutralizing mAbs that bind to the RBM that are not affected by Omicron mutations, including a molecular mimic of the ACE2 receptor (S2K146)^1^. Collectively, these data may guide future efforts to develop SARS-CoV-2 vaccines and therapies to counteract antigenic shift and future sarbecovirus zoonotic spillovers.

## Acknowledgements

We thank Hideki Tani (University of Toyama) for providing the reagents necessary for preparing VSV pseudotyped viruses. This study was supported by the National Institute of Allergy and Infectious Diseases (DP1AI158186 and HHSN272201700059C to D.V.), a Pew Biomedical Scholars Award (D.V.), an Investigators in the Pathogenesis of Infectious Disease Awards from the Burroughs Wellcome Fund (D.V.), Fast Grants (D.V.), the National Institute of General Medical Sciences (5T32GM008268-32 to SKZ). D.V. is an Investigator of the Howard Hughes Medical Institute. OG is funded by the Swiss Kidney Foundation. This work was supported, in part, by the National Institutes of Allergy and Infectious Diseases Center for Research on Influenza Pathogenesis (HHSN272201400008C), Center for Research on Influenza Pathogenesis and Transmission (CRIPT) (75N93021C00014), and the Japan Program for Infectious Diseases Research and Infrastructure (JP21wm0125002) from the Japan Agency for Medical Research and Development (AMED).

## Author contributions

Conceived research and designed study: D.C., G.S., M.S.P., L.P., D.V. Designed experiments: D.C., D.P., E.C., L.E.R., G.S., M.S.P., L.P., J.E.B., A.C.W., D.V. Designed and performed mutagenesis for S mutant expression plasmids: E.C. and K.C. Produced pseudoviruses: C.S., D.P., H.K., J.N., N.F., K.R.S. Carried out pseudovirus neutralization assays: C.S., J.E.B., D.P., F.Z., J.B., C.S-F. and A.D.M. C.S., K.C. and E.C. expressed antibodies. Isolation and propagation of SARS-CoV-2 Omicron live virus: L.A.V., P.J.H., Y.K. Carried out live virus neutralization assays: L.A.V., P.J.H. Supervised the research on live virus neutralization assays: M.S.D. L.E.R. performed binding assays. Cl.G., S.K.Z., A.C.W., N.C., A.E.P. and J.R.D. synthesized expression plasmid, expressed and purified ACE2 and RBD proteins. Production and quality control of mAbs: C.S., A.C. Bioinformatic and epidemiology analyses: J.diI., C.M., L.Y., D.S., L.S. Interpreted Data: C.S., D.P., L.P., L.E.R., M.S.P., A.D.M. Data analysis: E.C., C.S., F.Z., A.D.M., K.C., D.P., J.E.B., L.E.R., A.C.W., D.V., A.T., G.S., D.C. A.R., O.G., Ch.G., A.C., P.F., A.F.P., H.C., N.M.F., J.L., N.T.I., I.M., J.G., R.G., A.G, P.C. and C.H.D. contributed to donors recruitment and plasma samples collection. D.C., A.L., H.W.V., G.S., A.T., L.A.P., D.V., wrote the manuscript with input from all authors.

## MATERIALS AND METHODS

### Cell lines

Cell lines used in this study were obtained from ATCC (HEK293T and Vero E6), ThermoFisher Scientific (Expi CHO cells, FreeStyle™ 293-F cells and Expi293F™ cells) or generated in-house (Vero E6/TMPRSS2)^44^.

### Omicron prevalence analysis

The viral sequences and the corresponding metadata were obtained from GISAID EpiCoV project (https://www.gisaid.org/). Analysis was performed on sequences submitted to GISAID up to Dec 09, 2021. S protein sequences were either obtained directly from the protein dump provided by GISAID or, for the latest submitted sequences that were not incorporated yet in the protein dump at the day of data retrieval, from the genomic sequences with the exonerate^55^ 2 2.4.0-- haf93ef1_3 (https://quay.io/repository/biocontainers/exonerate?tab=tags) using protein to DNA alignment with parameters -m protein2dna --refine full --minintron 999999 --percent 20 and using accession YP_009724390.1 as a reference. Multiple sequence alignment of all human spike proteins was performed with mafft^56^ 7.475--h516909a_0 (https://quay.io/repository/biocontainers/mafft?tab=tags) with parameters --auto --reorder -- keeplength --addfragments using the same reference as above. S sequences that contained >10% ambiguous amino acid or that were < than 80% of the canonical protein length were discarded. Figures were generated with R 4.0.2 (https://cran.r-project.org/) using ggplot2 3.3.2 and sf 0.9-7 packages. To identify each mutation prevalence, missingness (or ambiguous amino acids) was taken into account in both nominator and denominator.

### Monoclonal Antibodies

VIR-7831 and VIR-7832 were produced at WuXi Biologics (China). Antibody VH and VL sequences for mAbs cilgavimab (PDB ID 7L7E), tixagevimab (PDB ID 7L7E, 7L7D), casirivimab (PDB ID 6XDG), imdevimab (PDB ID 6XDG) and ADI-58125 (PCT application WO2021207597, seq. IDs 22301 and 22311) were subcloned into heavy chain (human IgG1) and the corresponding light chain (human IgKappa, IgLambda) expression vectors respectively and produced in transiently expressed in Expi-CHO-S cells (Thermo Fisher, #A29133) at 37°C and 8% CO2. Cells were transfected using ExpiFectamine. Transfected cells were supplemented 1 day after transfection with ExpiCHO Feed and ExpiFectamine CHO Enhancer. Cell culture supernatant was collected eight days after transfection and filtered through a 0.2 μm filter. Recombinant antibodies were affinity purified on an ÄKTA Xpress FPLC device using 5 mL HiTrap™ MabSelect™ PrismA columns followed by buffer exchange to Histidine buffer (20 mM Histidine, 8% sucrose, pH 6) using HiPrep 26/10 desalting columns. Antibody VH and VL sequences for bamlanivimab (LY-CoV555), etesevimab (LY-CoV016), regdanvimab (CT-P59) were obtained from PDB IDs 7KMG, 7C01 and 7CM4, respectively and mAbs were produced as recombinant IgG1 by ATUM. The mAbs composing the NTD- and RBD-specific were discovered at VIR Biotechnology and have been produced as recombinant IgG1 in Expi-CHO-S cells as described above. The identity of the produced mAbs was confirmed by LC-MS analysis.

### IgG mass quantification by LC/MS intact protein mass analysis

Fc N-linked glycan from mAbs were removed by PNGase F after overnight non-denaturing reaction at room temperature. Deglycosylated protein (4 μg) was injected to the LC-MS system to acquire intact MS signal. Thermo MS (Q Exactive Plus Orbitrap) was used to acquire intact protein mass under denaturing condition with m/z window from 1,000 to 6,000. BioPharma Finder 3.2 software was used to deconvolute the raw m/z data to protein average mass. The theoretical mass for each mAb was calculated with GPMAW 10.10 software. Many of the protein post-translational modifications such as N-terminal pyroglutamate cyclization, and c-terminal lysine cleavage, and formation of 16-18 disulfide bonds were added into the calculation.

### Sample donors

Samples were obtained from SARS-CoV-2 recovered and vaccinated individuals under study protocols approved by the local Institutional Review Boards (Canton Ticino Ethics Committee, Switzerland, Comitato Etico Milano Area 1). All donors provided written informed consent for the use of blood and blood derivatives (such as PBMCs, sera or plasma) for research. Samples were collected 14-28 days after symptoms onset and 14-28 days or 7-10 months after vaccination. Convalescent plasma, Ad26.COV2.S, mRNA-1273 and BNT162b2 samples were obtained from the HAARVI study approved by the University of Washington Human Subjects Division Institutional Review Board (STUDY00000959). AZD1222 samples were obtained from INGM, Ospedale Maggio Policlinico of Milan and approved by the local review board Study Polimmune. Sputnik V samples were obtained from healthcare workers at the hospital de Clínicas “José de San Martín”, Buenos Aires, Argentina. Sinopharm vaccinated individuals were enrolled from Aga Khan University under IRB of UWARN study.

### Serum/plasma and mAbs pseudovirus neutralization assays

#### VSV pseudovirus generation used on Vero E6 cells

The plasmids encoding the Omicron SARS-CoV-2 S variant was generated by overlap PCR mutagenesis of the wild-type plasmid, pcDNA3.1(+)-spike-D19^57^. Replication defective VSV pseudovirus expressing SARS-CoV-2 spike proteins corresponding to the ancestral Wuhan-Hu-1 virus and the Omicron VOC were generated as previously described^8^ with some modifications. Lenti-X 293T cells (Takara) were seeded in 15-cm^2^ dishes at a density of 10e6 cells per dish and the following day transfected with 25 μg of spike expression plasmid with TransIT-Lenti (Mirus, 6600) according to the manufacturer’s instructions. One day post-transfection, cells were infected with VSV-luc (VSV-G) with an MOI 3 for 1 h, rinsed three times with PBS containing Ca2+/Mg2+, then incubated for additional 24 h in complete media at 37°C. The cell supernatant was clarified by centrifugation, aliquoted, and frozen at -80°C.

#### VSV pseudovirus generation used on Vero E6-TMPRSS2 cells

Comparison of Omicron SARS-CoV-2 S VSV to SARS-CoV-2 G614 S (YP 009724390.1) VSV and Beta S VSV used pseudotyped particles prepared as described previously^10,58^. Briefly, HEK293T cells in DMEM supplemented with 10% FBS, 1% PenStrep seeded in 10-cm dishes were transfected with the plasmid encoding for the corresponding S glycoprotein using lipofectamine 2000 (Life Technologies) following the manufacturer’s instructions. One day post-transfection, cells were infected with VSV(G*ΔG-luciferase)^59^ and after 2 h were washed five times with DMEM before adding medium supplemented with anti-VSV-G antibody (I1-mouse hybridoma supernatant, CRL-2700, ATCC). Virus pseudotypes were harvested 18-24 h post-inoculation, clarified by centrifugation at 2,500 x g for 5 min, filtered through a 0.45 μm cut off membrane, concentrated 10 times with a 30 kDa cut off membrane, aliquoted and stored at -80°C.

### VSV pseudovirus neutralization

#### Assay performed using Vero E6 cells

Vero-E6 were grown in DMEM supplemented with 10% FBS and seeded into clear bottom white 96 well plates (PerkinElmer, 6005688) at a density of 20’000 cells per well. The next day, mAbs or plasma were serially diluted in pre-warmed complete media, mixed with pseudoviruses and incubated for 1 h at 37°C in round bottom polypropylene plates. Media from cells was aspirated and 50 μl of virus-mAb/plasma complexes were added to cells and then incubated for 1 h at 37°C. An additional 100 μL of prewarmed complete media was then added on top of complexes and cells incubated for an additional 16-24 h. Conditions were tested in duplicate wells on each plate and eight wells per plate contained untreated infected cells (defining the 0% of neutralization, “MAX RLU” value) and infected cells in the presence of S309 and S2X259 at 20 μg/ml each (defining the 100% of neutralization, “MIN RLU” value). Virus-mAb/plasma-containing media was then aspirated from cells and 100 μL of a 1:2 dilution of SteadyLite Plus (Perkin Elmer, 6066759) in PBS with Ca^++^ and Mg^++^ was added to cells. Plates were incubated for 15 min at room temperature and then were analyzed on the Synergy-H1 (Biotek). Average of Relative light units (RLUs) of untreated infected wells (MAX RLU_ave_) was subtracted by the average of MIN RLU (MIN RLU_ave_) and used to normalize percentage of neutralization of individual RLU values of experimental data according to the following formula: (1-(RLU_x_ -MIN RLU_ave_) / (MAX RLU_ave_ – MIN RLU_ave_)) x 100. Data were analyzed and visualized with Prism (Version 9.1.0). IC50 (mAbs) and ID50 (plasma) values were calculated from the interpolated value from the log(inhibitor) versus response, using variable slope (four parameters) nonlinear regression with an upper constraint of ≤100, and a lower constrain equal to 0. Each neutralization experiment was conducted on two independent experiments, i.e., biological replicates, where each biological replicate contains a technical duplicate. IC50 values across biological replicates are presented as arithmetic mean ± standard deviation. The loss or gain of neutralization potency across spike variants was calculated by dividing the variant IC50/ID50 by the parental IC50/ID50 within each biological replicate, and then visualized as arithmetic mean ± standard deviation.

#### Assay performed using Vero E6-TMPRSS2 cells

VeroE6-TMPRSS2 were cultured in DMEM with 10% FBS (Hyclone), 1% PenStrep and 8 μg/mL puromycin (to ensure retention of TMPRSS2) with 5% CO_2_ in a 37°C incubator (ThermoFisher). Cells were trypsinized using 0.05% trypsin and plated to be at 90% confluence the following day. In an empty half-area 96-well plate, a 1:3 serial dilution of sera was made in DMEM and diluted pseudovirus was then added and incubated at room temperature for 30-60 min before addition of the sera-virus mixture to the cells at 37°C. 2 hours later, 40 μL of a DMEM solution containing 20% FBS and 2% PenStrep was added to each well. After 17-20 hours, 40 μL/well of One-Glo-EX substrate (Promega) was added to the cells and incubated in the dark for 5-10 min prior to reading on a BioTek plate reader. Measurements were done at least in duplicate using distinct batches of pseudoviruses and one representative experiment is shown. Relative luciferase units were plotted and normalized in Prism (GraphPad). Nonlinear regression of log(inhibitor) versus normalized response was used to determine IC_50_ values from curve fits. Normality was tested using the D’Agostino-Pearson test and in the absence of a normal distribution, Kruskal-Wallis tests were used to compare two groups to determine whether differences reached statistical significance. Fold changes were determined by comparing individual IC_50_ and then averaging the individual fold changes for reporting.

### Focus reduction neutralization test

Vero-TMPRSS2^60^ cells were cultured at 37°C in Dulbecco’s Modified Eagle medium (DMEM) supplemented with 10% fetal bovine serum (FBS), 10 mM HEPES pH 7.3, and 100 U/ml of penicillin– streptomycin and supplemented with 5 μg/mL of blasticidin. The WA1/2020 strain with a D614G substitution was described previously^61^. The B.1.1.529 isolate (hCoV-19/USA/WI-WSLH-221686/2021) was obtained from a nasal swab and passaged on Vero-TMPRSS2 cells as described^62^. The B.1.1.529 isolate was sequenced (GISAID: EPI_ISL_7263803) to confirm the stability of substitutions. All virus experiments were performed in an approved biosafety level 3 (BSL-3) facility.

Serial dilutions of VIR-7381 mAbs were incubated with 10^2^ focus-forming units (FFU) of SARS-CoV-2 (WA1/2020 D614G or B.1.1.529) for 1 h at 37°C. Antibody-virus complexes were added to Vero-TMPRSS2 cell monolayers in 96-well plates and incubated at 37°C for 1 h. Subsequently, cells were overlaid with 1% (w/v) methylcellulose in MEM. Plates were harvested at 30 h (WA1/2020 D614G on Vero-TMPRSS2 cells) or 70 h (B.1.1.529 on Vero-TMPRSS2 cells) later by removal of overlays and fixation with 4% PFA in PBS for 20 min at room temperature. Plates with WA1/2020 D614G were washed and sequentially incubated with an oligoclonal pool of SARS2-2, SARS2-11, SARS2-16, SARS2-31, SARS2-38, SARS2-57, and SARS2-71^63^ anti-S antibodies. Plates with B.1.1.529 were additionally incubated with a pool of mAbs that cross-react with SARS-CoV-1 and bind a CR3022-competing epitope on the RBD^64^. All plates were subsequently stained with HRP-conjugated goat anti-mouse IgG (Sigma, A8924) in PBS supplemented with 0.1% saponin and 0.1% bovine serum albumin. SARS-CoV-2-infected cell foci were visualized using TrueBlue peroxidase substrate (KPL) and quantitated on an ImmunoSpot microanalyzer (Cellular Technologies). Antibody-dose response curves were analyzed using non-linear regression analysis with a variable slope (GraphPad Software), and the half-maximal inhibitory concentration (IC_50_) was calculated.

### Recombinant RBD and hACE2 protein production

SARS-CoV-2 RBD proteins for SPR binding assays (residues 328-531from GenBank for WT: NC_045512.2 with N-terminal signal peptide and C-terminal thrombin cleavage site-TwinStrep-8xHis-tag) were expressed in Expi293F (Thermo Fisher Scientific) cells at 37°C and 8% CO2. Transfections were performed using the ExpiFectamine 293 Transfection Kit (Thermo Fisher Scientific) using RBD expression plasmids produced at ATUM. Cell culture supernatants were collected two to four days after transfection and supplemented with 10x PBS to a final concentration of 2.5x PBS (342.5 mM NaCl, 6.75 mM KCl and 29.75 mM phosphates). SARS-CoV-2 RBDs were purified using cobalt-based immobilized metal affinity chromatography followed by buffer exchange into PBS using a HiPrep 26/10 desalting column (Cytiva) or a Superdex 200 Increase 10/300 GL column (Cytiva), for the two batches of Omicron RBD used for SPR, respectively. Recombinant human ACE2 (residues 19-615 from Uniprot Q9BYF1 with a C-terminal AviTag-10xHis-GGG-tag, and N-terminal signal peptide) was produced by ATUM. Protein was purified via Ni Sepharose resin followed by isolation of the monomeric hACE2 by size exclusion chromatography using a Superdex 200 Increase 10/300 GL column (Cytiva) pre-equilibrated with PBS.

### Transient expression and purification of animal ACE2

The mouse (GenBank: Q8R0I0), american mink (GenBank: QPL12211.1), and pangolin (XP_017505752.1) ACE2 ectodomains constructs were synthesized by GenScript and placed into a pCMV plasmid. The domain boundaries for the ectodomain are residues 19-615. The native signal tag was identified using SignalP-5.0 (residues 1-18) and replaced with a N-terminal mu-phosphatase signal peptide. These constructs were then fused to a sequence encoding thrombin cleavage site and a human Fc fragment or a 8x His tag at the C-terminus. All ACE2-Fc, and ACE2 His constructs were produced in Expi293 cells (Thermo Fisher A14527) in Gibco Expi293 Expression Medium at 37°C in a humidified 8% CO2 incubator rotating at 130 rpm. The cultures were transfected using PEI-25K (Polyscience) with cells grown to a density of 3 million cells per mL and cultivated for 4-5 days. Proteins were purified from clarified supernatants for using a 1 mL HiTrap Protein A HP affinity column (Cytiva) or a 1 mL HisTrap HP affinity column (Cytiva), concentrated and flash frozen in 1x PBS, pH 7.4 (10 mM Na2HPO4, 1.8 mM KH2PO4, 2.7 mM KCl, 137 mM NaCl).

### ACE2 binding measurements using surface plasmon resonance

Measurements were performed using a Biacore T200 instrument, in triplicate for monomeric human and mouse ACE2 and duplicate for dimeric animal ACE2. A CM5 chip covalently immobilized with StrepTactin XT was used for surface capture of StrepTag-containing RBDs. Two different batches of Omicron RBD were used for the experiments. Running buffer was HBS-EP+ pH 7.4 (Cytiva) and measurements were performed at 25 °C. Experiments were performed with a 3-fold dilution series of monomeric human ACE2 (300, 100, 33, 11 nM) or animal ACE2 (900, 300, 100, 33 nM) and were run as single-cycle kinetics. Data were double reference-subtracted and fit to a 1:1 binding model using Biacore Evaluation software.

### Statistical analysis

Neutralization measurements were done in duplicate and relative luciferase units were converted to percent neutralization and plotted with a non-linear regression model to determine IC50/ID50 values using GraphPad PRISM software (version 9.0.0). Comparisons between two groups of paired data were made with Wilcoxon rank test. Comparisons between multiple groups of unpaired data were made with Kruskal-Wallis rank test and corrected with Dunn’s test.

---

**Extended Data Table 1.** Characteristics of single point mutations present in Omicron RBD relative to Wuhan-Hu-1 RBD.

|  | **RBD expr.** | **ACE2 binding** | **ACE2 binding** |
| --- | --- | --- | --- |
|  |  | FACS | SPR |
|  | log fold-change | log fold-change | fold-change |
| **G339D** | 0.3 | 0.06 |  |
| **S371L** | -0.61 | -0.14 |  |
| **S373P** | -0.22 | -0.08 |  |
| **S375F** | -1.81 | -0.55 |  |
| **K417N** | 0.1 | -0.45 | 0.3 |
| **N440K** | -0.12 | 0.07 | 1.2 |
| **G446S** | -0.4 | -0.2 |  |
| **S477N** | 0.06 | 0.06 |  |
| **T478K** | 0.02 | 0.02 | 1.2 |
| **E484A** | -0.23 | -0.07 | 1.0 for E484K |
| **Q493K** | 0.03 | 0.05 |  |
| **G496S** | 0.12 | -0.63 |  |
| **Q498R** | -0.1 | -0.06 |  |
| **N501Y** | -0.14 | 0.24 | 6.2 |
| **Y505H** | 0.16 | -0.71 |  |
| **Total:** | -2.84 | -2.39 |  |
|  | (692-fold loss) | (245-fold loss) |  |
| **K417N/**<br>**E484K/**<br>**N501Y** |  |  | 2.4 |

Expression and FACS ACE2 binding data from<sup>26</sup>

**Extended Data Table 2.**
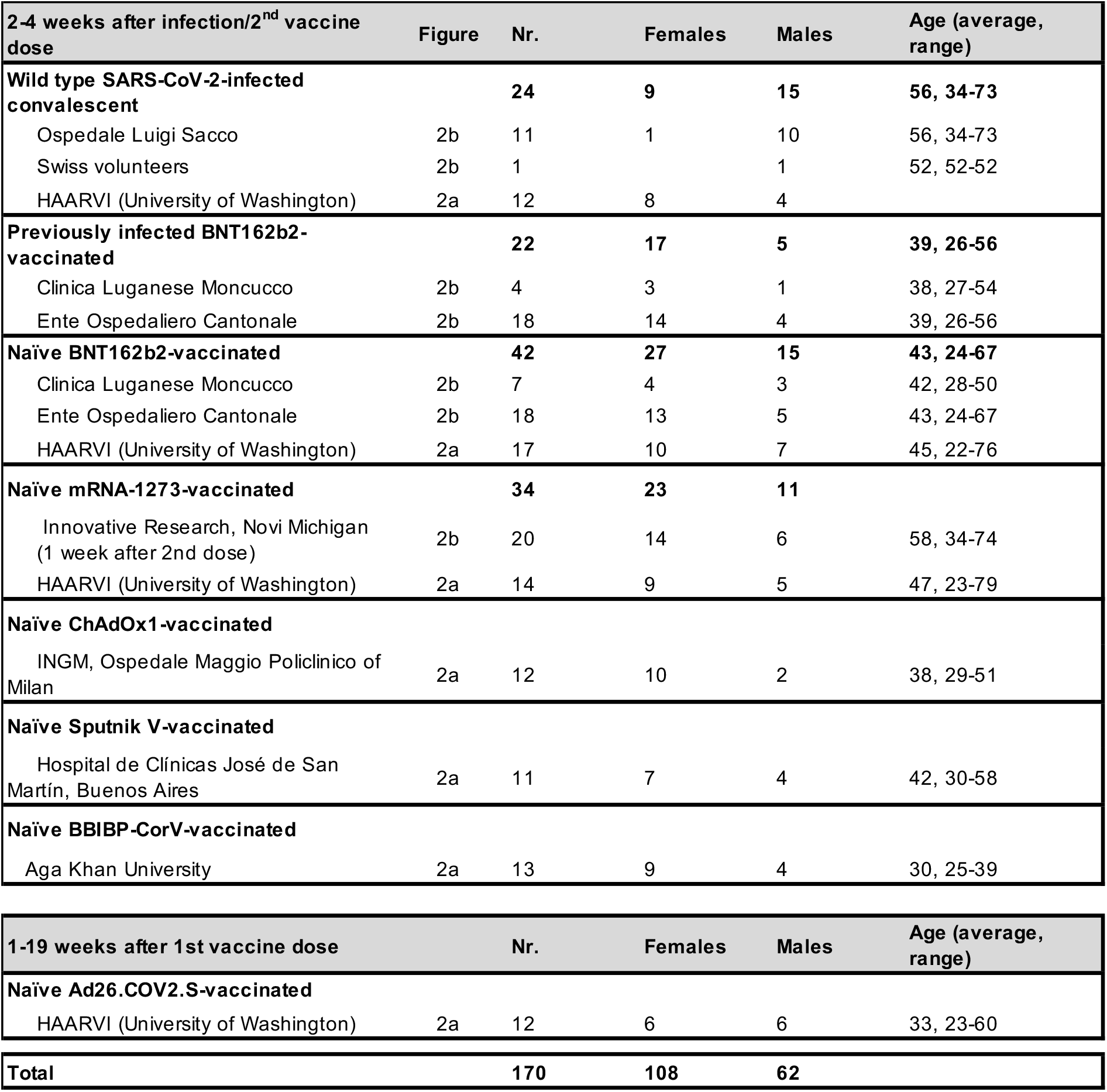
Enrolled donors’ demographics.

**Extended Data Table 3.**
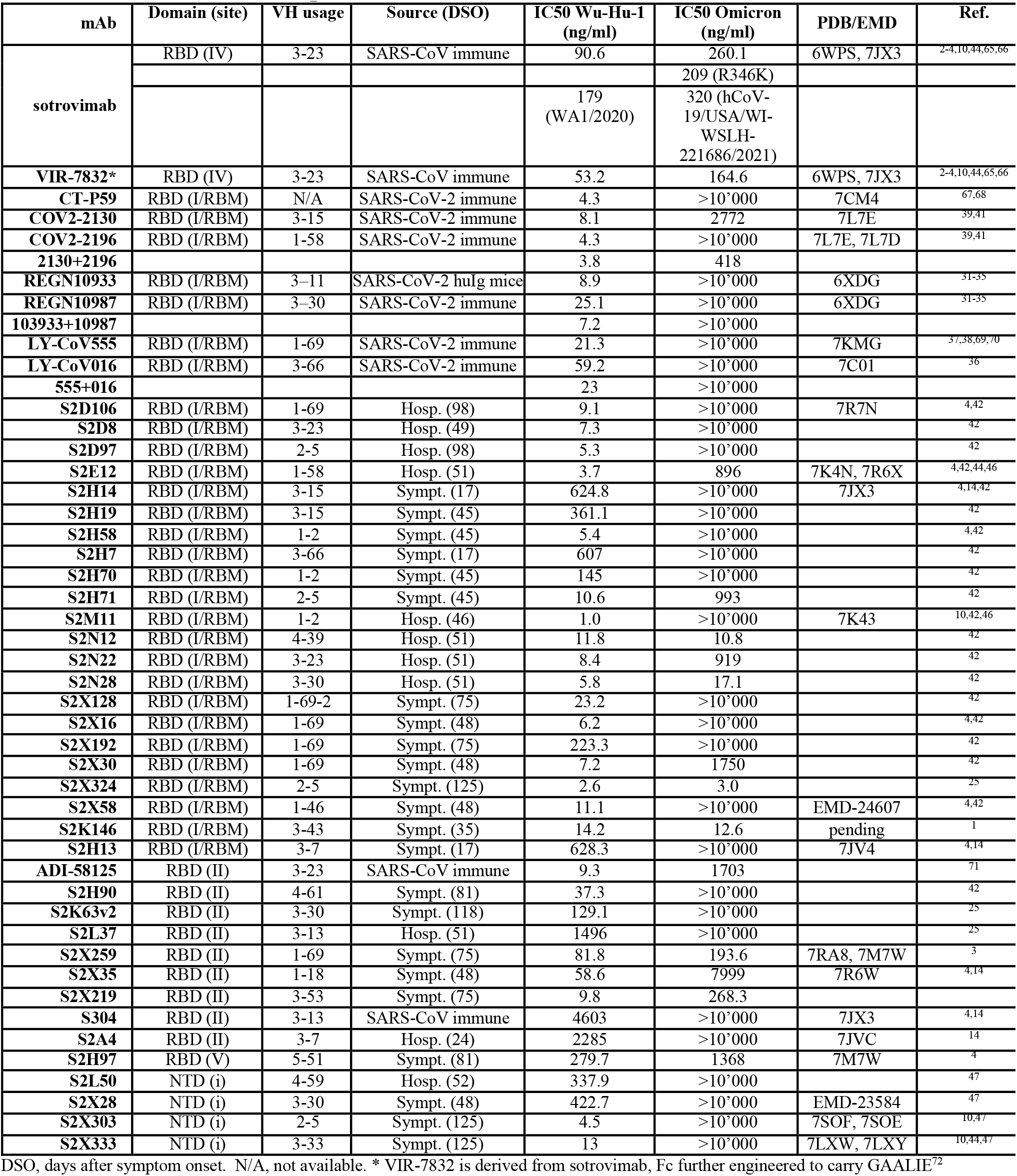
Properties of tested mAbs.

**Extended Data Fig. 1.**
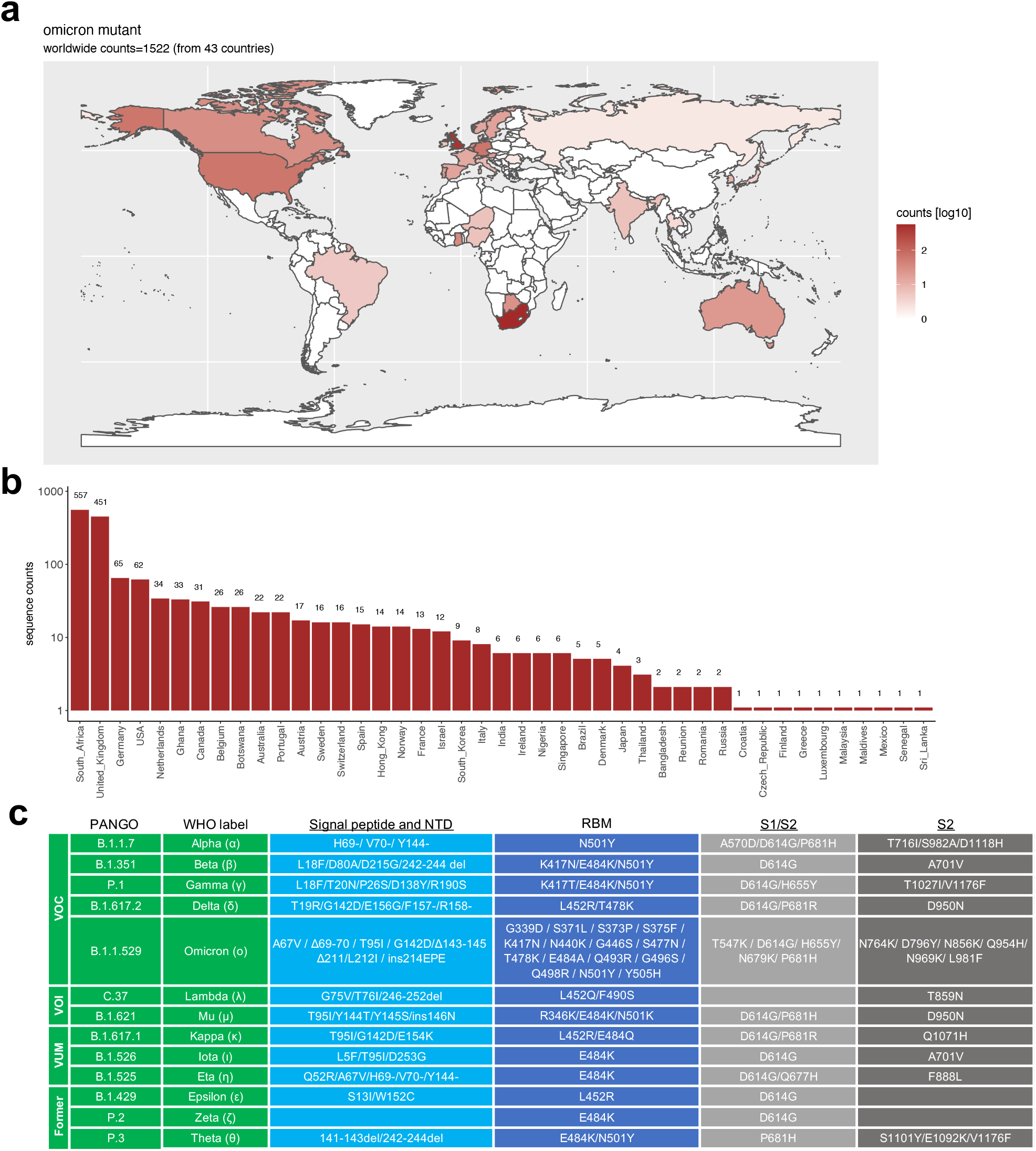
Geographic distribution and evolution of incidence over time of the SARS-COV-2 Omicron VOC and schematic of mutations landscape. **a**, World map showing the geographic distribution and sequence counts of Omicron as of December 9, 2021. **b**, Total number of Omicron sequences deposited by country as of December 9, 2021. **c**, Fraction (left) and total number (right) of sequences deposited on a weekly basis worldwide (grey) or in South Africa (red). **c**, Schematic of mutations landscape in each current and former SARS-CoV-2 VOC, VOI and VUM (Variant Under Monitoring). b, deletion: ins, insertion.

**Extended Data Fig. 2.**
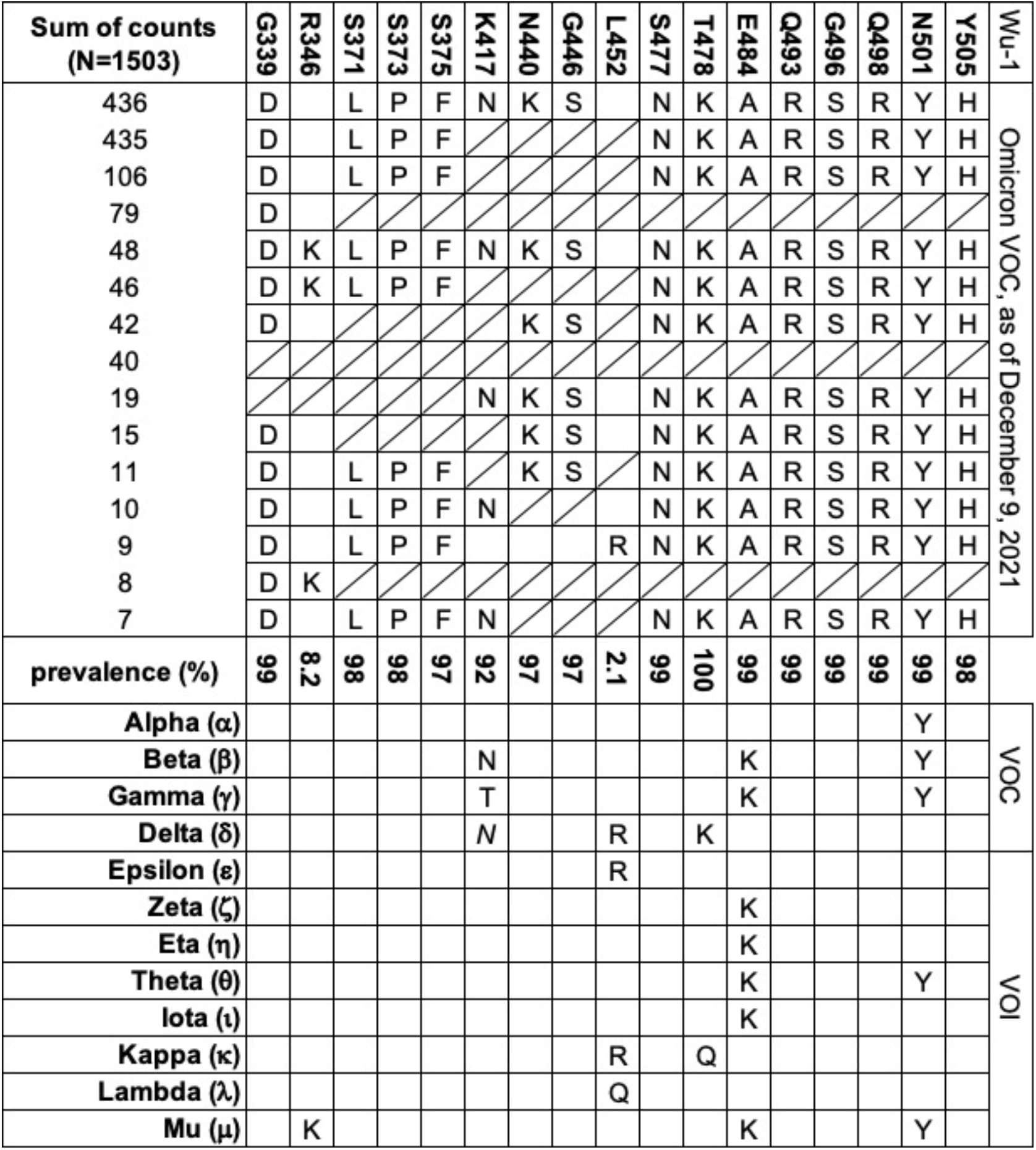
Amino acid substitutions and their prevalence in the Omicron RBD sequences reported in GIAID as of December 9, 2021; (ambiguous amino acid substitutions are indicated with strikethrough cells). Shown are also the substitutions found in other variants. K417N mutation in Delta is found only in a fraction of sequences. K417N mutation in Delta is found only in a fraction of sequences.

**Extended Data Fig. 3.**
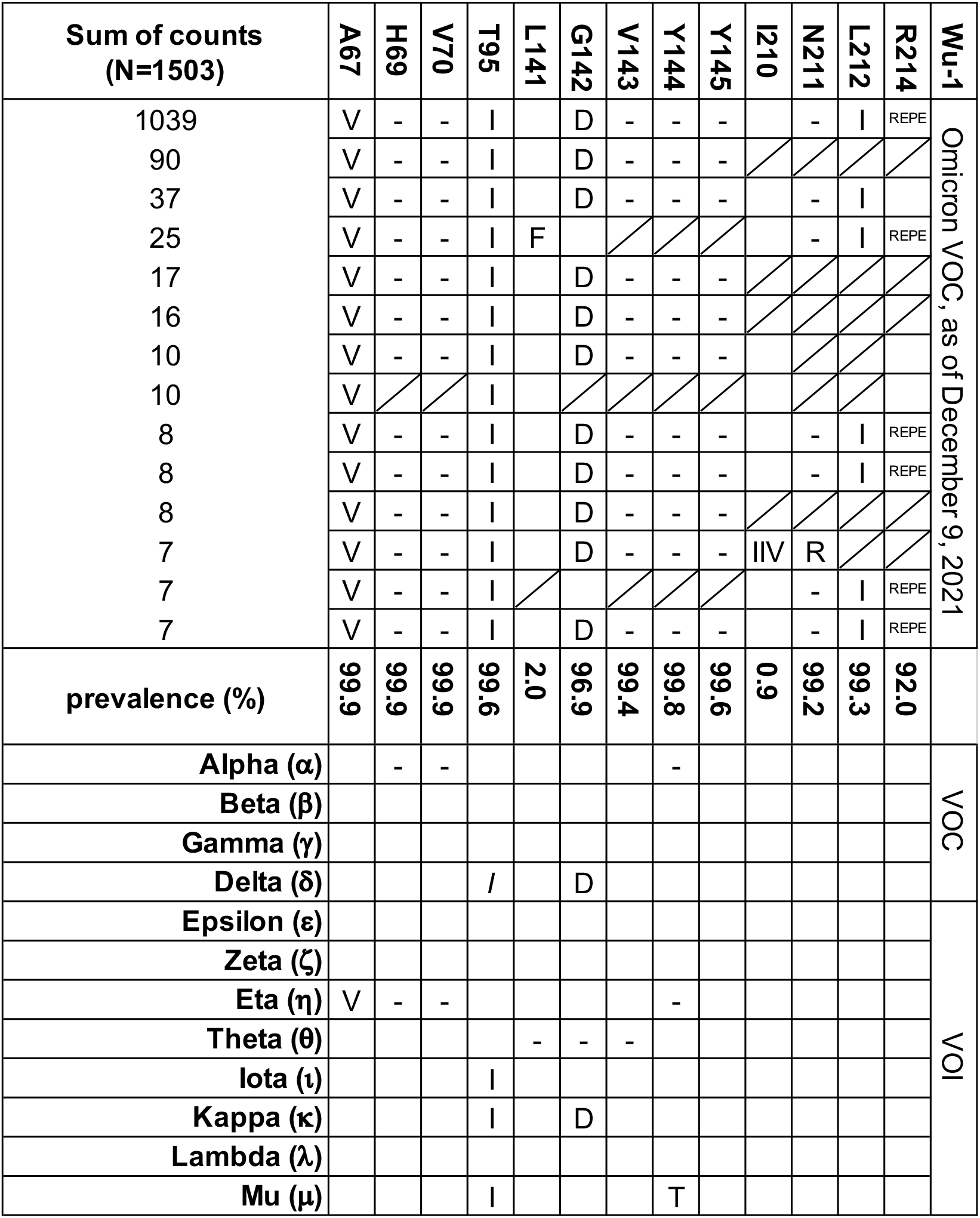
Amino acid substitutions and their prevalence in the Omicron NTD sequences reported in GIAID as of December 9, 2021; (ambiguous amino acid substitutions are marked with strikethrough cells). Shown are also the substitutions found in other variants.

**Extended Data Fig. 4.**
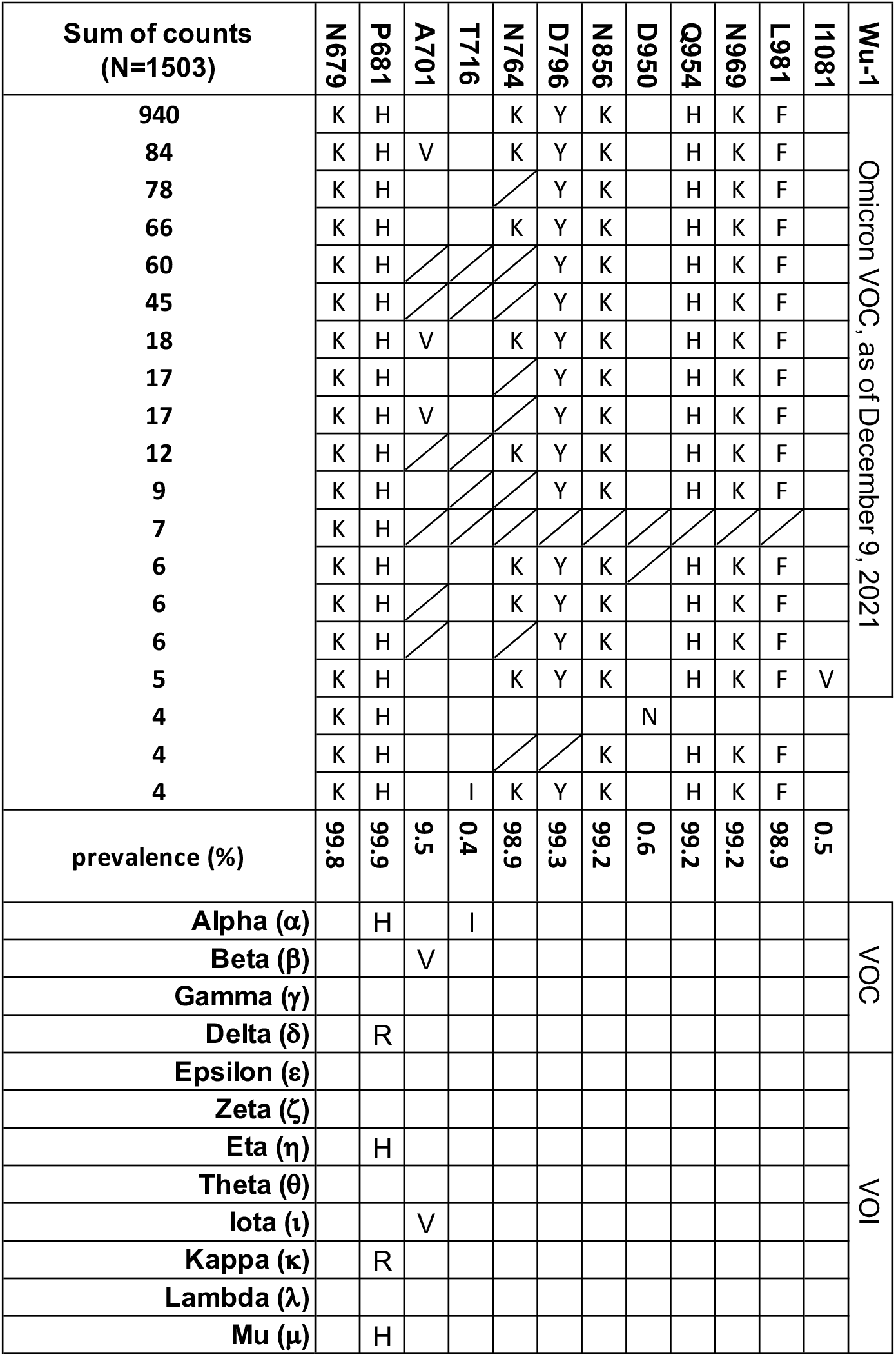
Amino acid substitutions and their prevalence in the Omicron S2 sequences reported in GIAID as of December 9, 2021; (ambiguous amino acid substitutions are marked with strikethrough cells). Shown are also the substitutions found in other variants.

**Extended Data Fig. 5.**
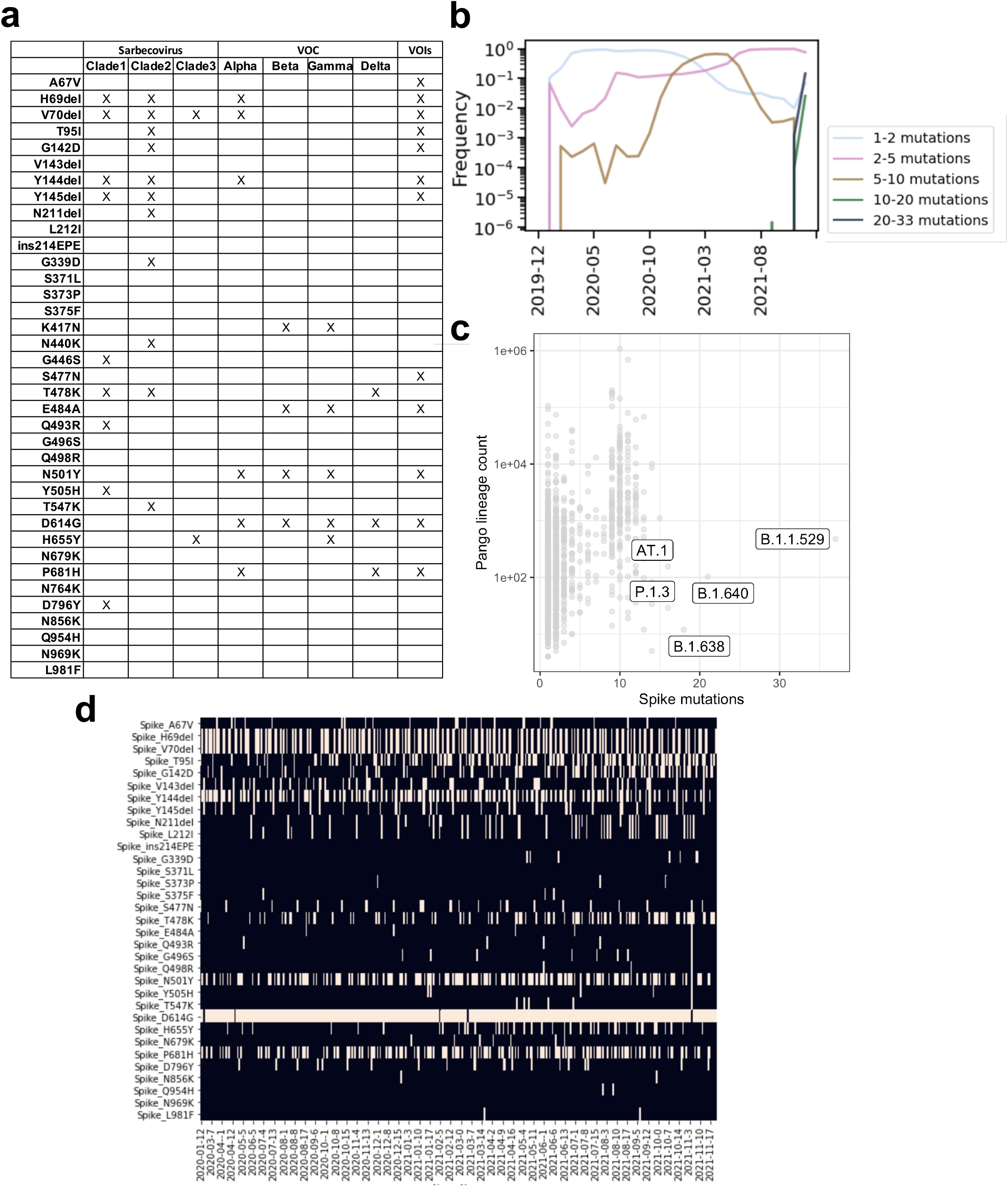
Characteristics of emergent mutations of Omicron. **a**, Shared mutations of micron with other sarbecovirus and with VOCs. **b**, Since the beginning of the pandemic there is a progressive coalescence of Omicron-defining mutations into non-Omicron haplotypes that may carry as many as 10 of the Omicron-defining mutations. **c**, Pango lineages (dots) rarely carry more than 10-15 lineage-defining mutations. **d**, Exceptionally, some non-Omicron haplotypes may carry up to a maximum 19 Omicron-defining mutations. Shown are selected exceptional haplotypes. Spike G142D and Y145del may also be noted as G142del and Y145D.

**Extended Data Fig. 6.**
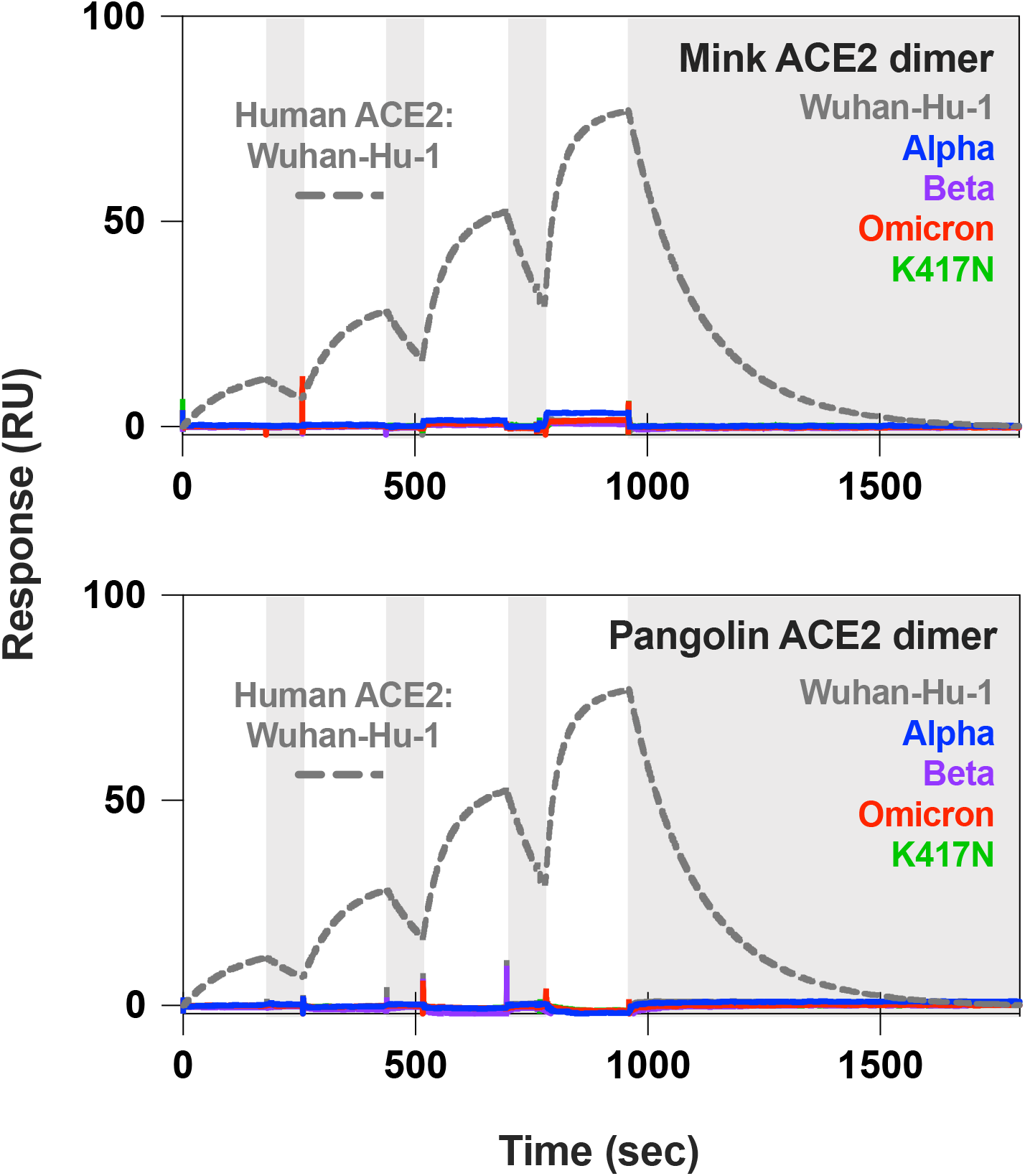
SPR analysis of pangolin and mink ACE2. Single-cycle kinetics SPR analysis of ACE2 binding to five RBD variants. Dimeric mink or pangolin ACE2 is injected successively at 33, 100, 300, and 900 nM. White and gray stripes indicate association and dissociation phases, respectively. Monomeric human ACE2 binding to Wuhan-Hu-1 RBD (ACE2 concentrations of 11, 33, 100, and 300 nM) shown for comparison.

**Extended Data Fig. 7.**
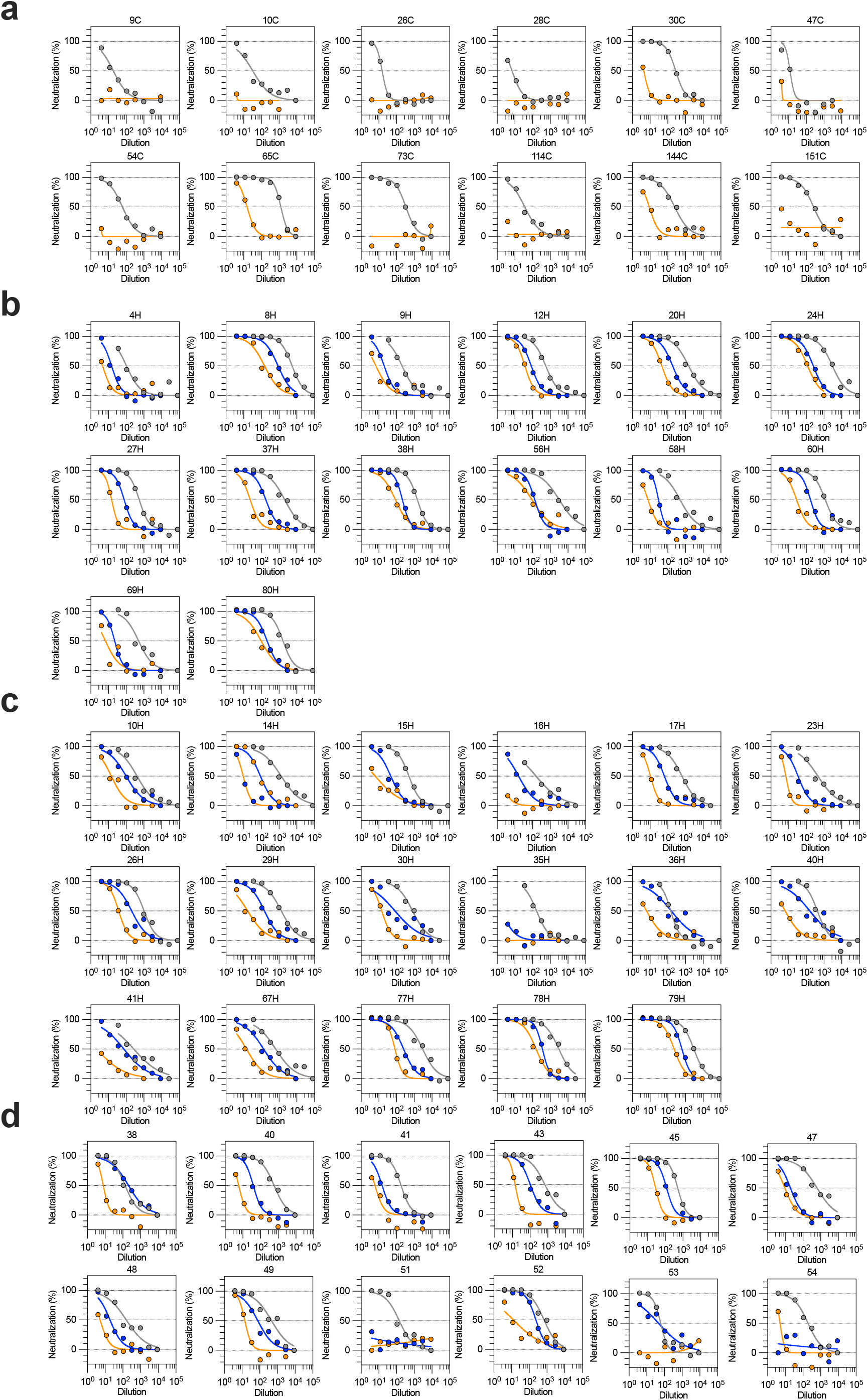
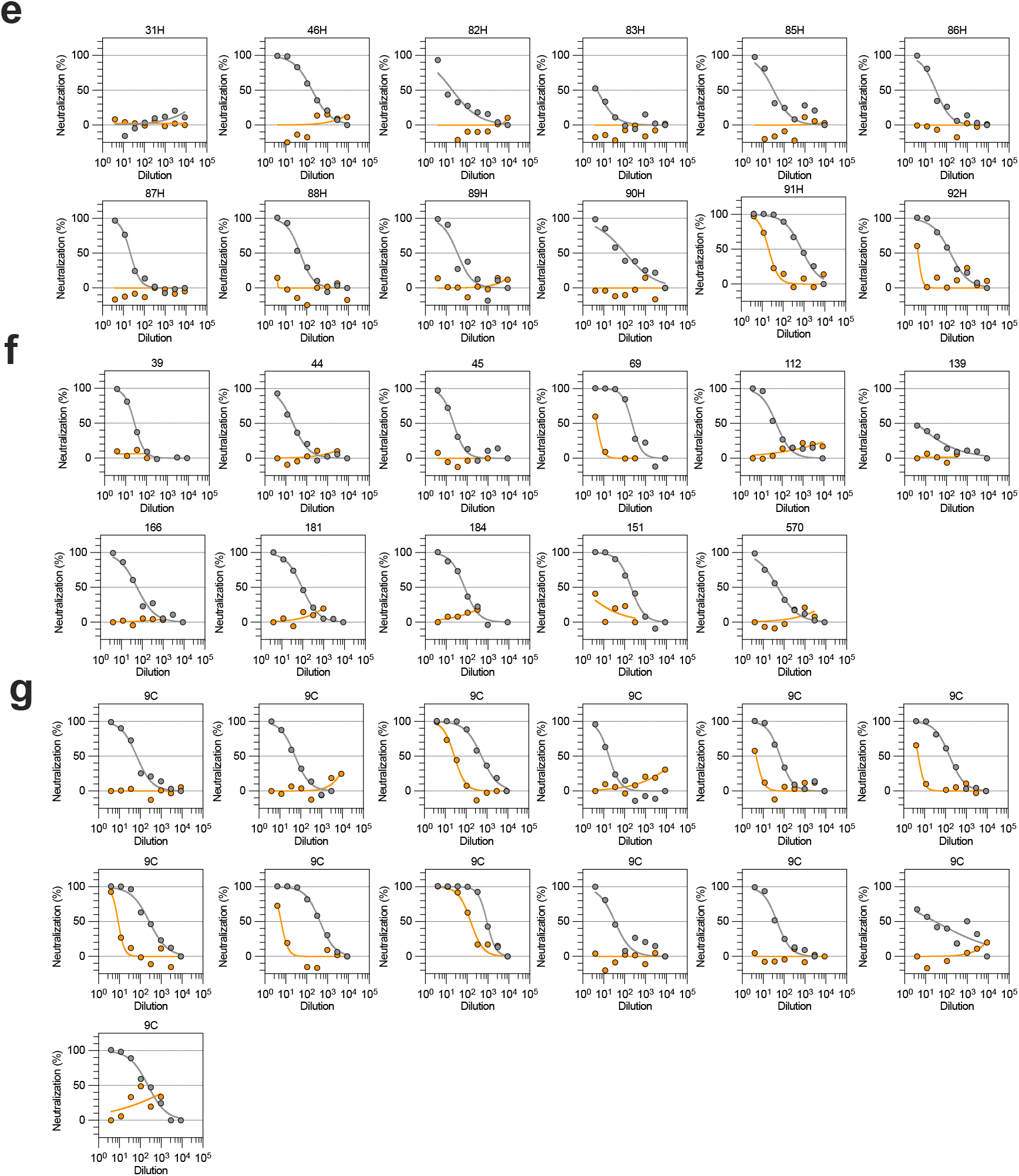
Neutralization of WT and Omicron S pseudotyped SARS-CoV-2 virus neutralization by plasma from COVID-19 convalescent and vaccinated individuals. Neutralization of SARS-CoV-2 pseudotyped VSV carrying Wuhan-Hu-1 D614G (grey), Beta (blue) or Omicron (orange) S protein by plasma from convalescent (**a**) or vaccinated individuals (**b**, mRNA-1273; **c**, BNT162b2; **d**, ChAdOx1; **e**, Ad26.COV2.S; **f**, Sputnik V; **g**, BBIBP-CorV) as shown in **Fig. 2a**. Data are representative of *n = 2* independent experiments.

**Extended Data Fig. 8.**
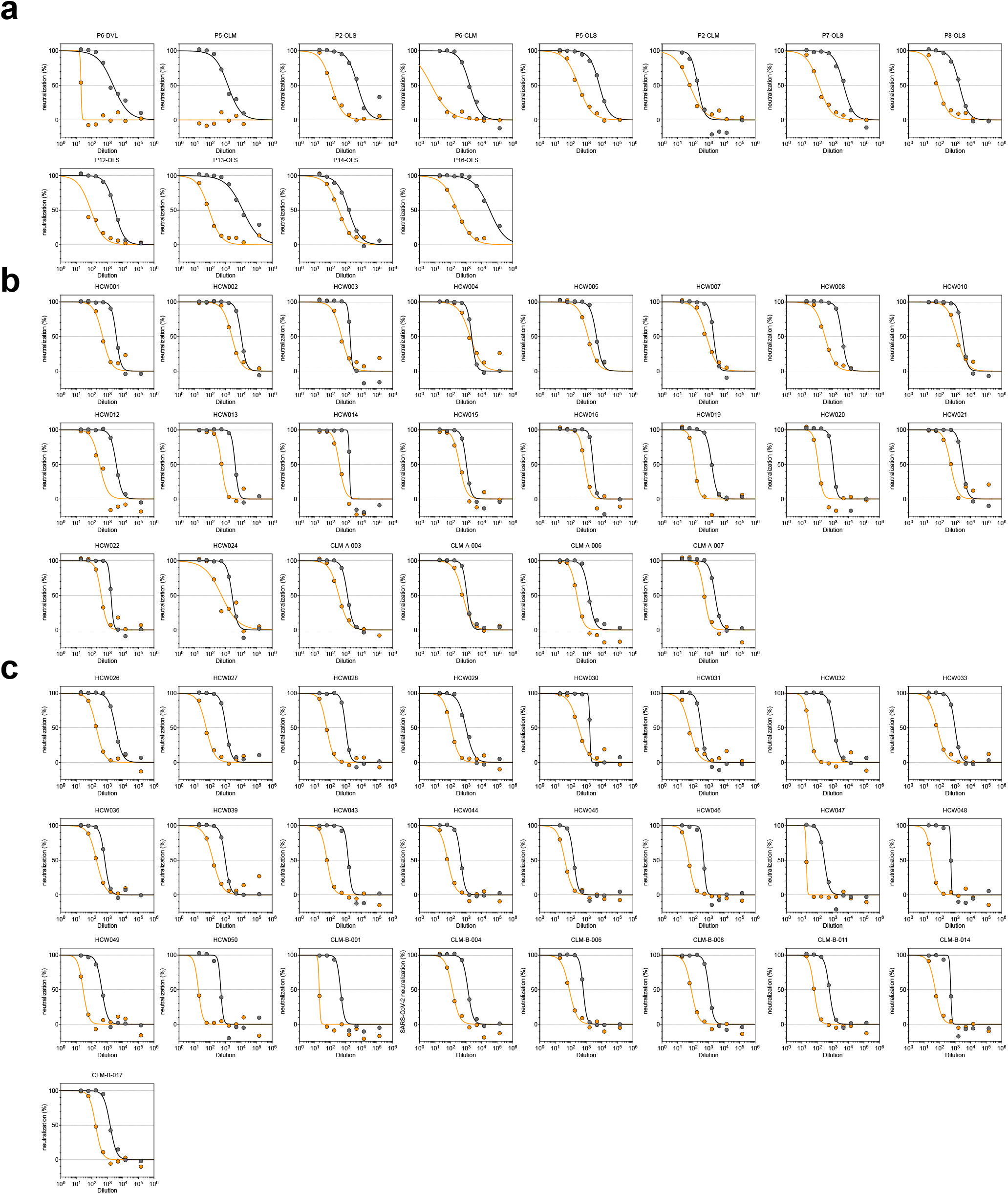
Neutralization of WT and Omicron S pseudotyped SARS-CoV-2 virus neutralization by plasma from COVID-19 convalescent and vaccinated individuals. **a-c**, Neutralization of SARS-CoV-2 pseudotyped VSV carrying Wuhan-Hu-1 or Omicron S protein by plasma from convalescent individuals 2-4 weeks after infection by WT SARS-CoV-2 (**a**, 11 out 12 individuals were hospitalized for COVID-19), and previously infected (**b**) or naïve (**c**) individuals, 2-4 weeks after receiving the second dose of BNT162b2 mRNA vaccine. Data are representative of *n = 2* independent experiments.

**Extended Data Fig. 9.**
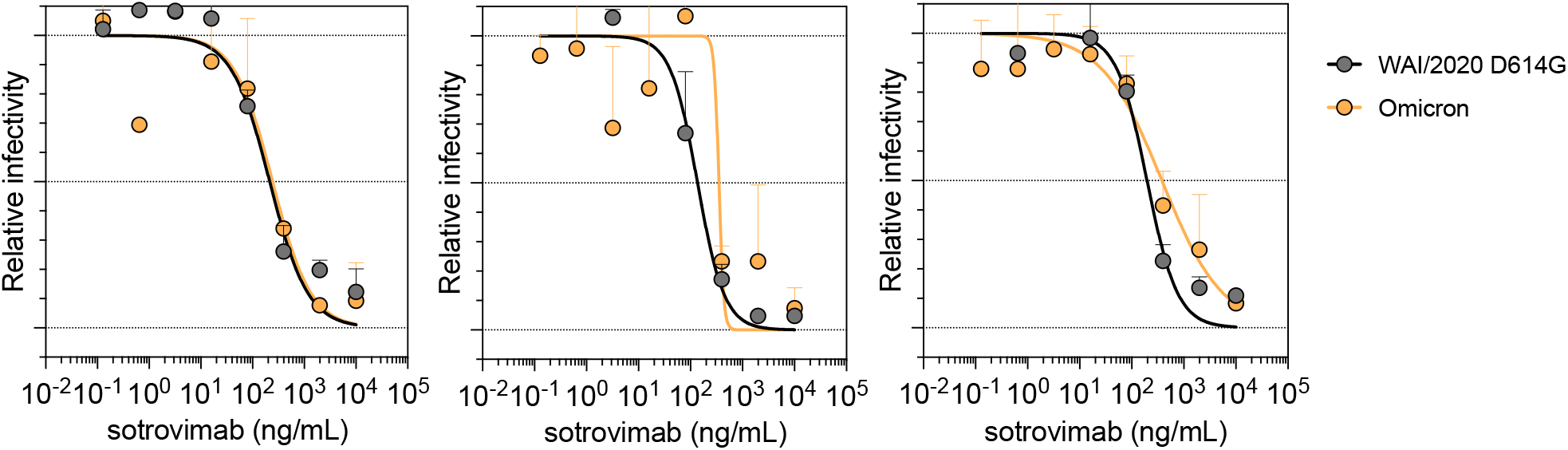
Neutralization of SARS-CoV-2 Omicron strain by sotrovimab in Vero-TMPRSS2 cells. **a-f**, Neutralization curves in Vero-TMPRSS2 cells comparing the sensitivity of SARS-CoV-2 strains with sotrovimab with WA1/2020 D614G and hCoV-19/USA/WI-WSLH-221686/2021 (an infectious clinical isolate of Omicron from a symptomatic individual in the United States). Shown are three independent experiments performed in technical duplicate is shown. Error bars indicate range.

**Extended Data Fig. 10.**
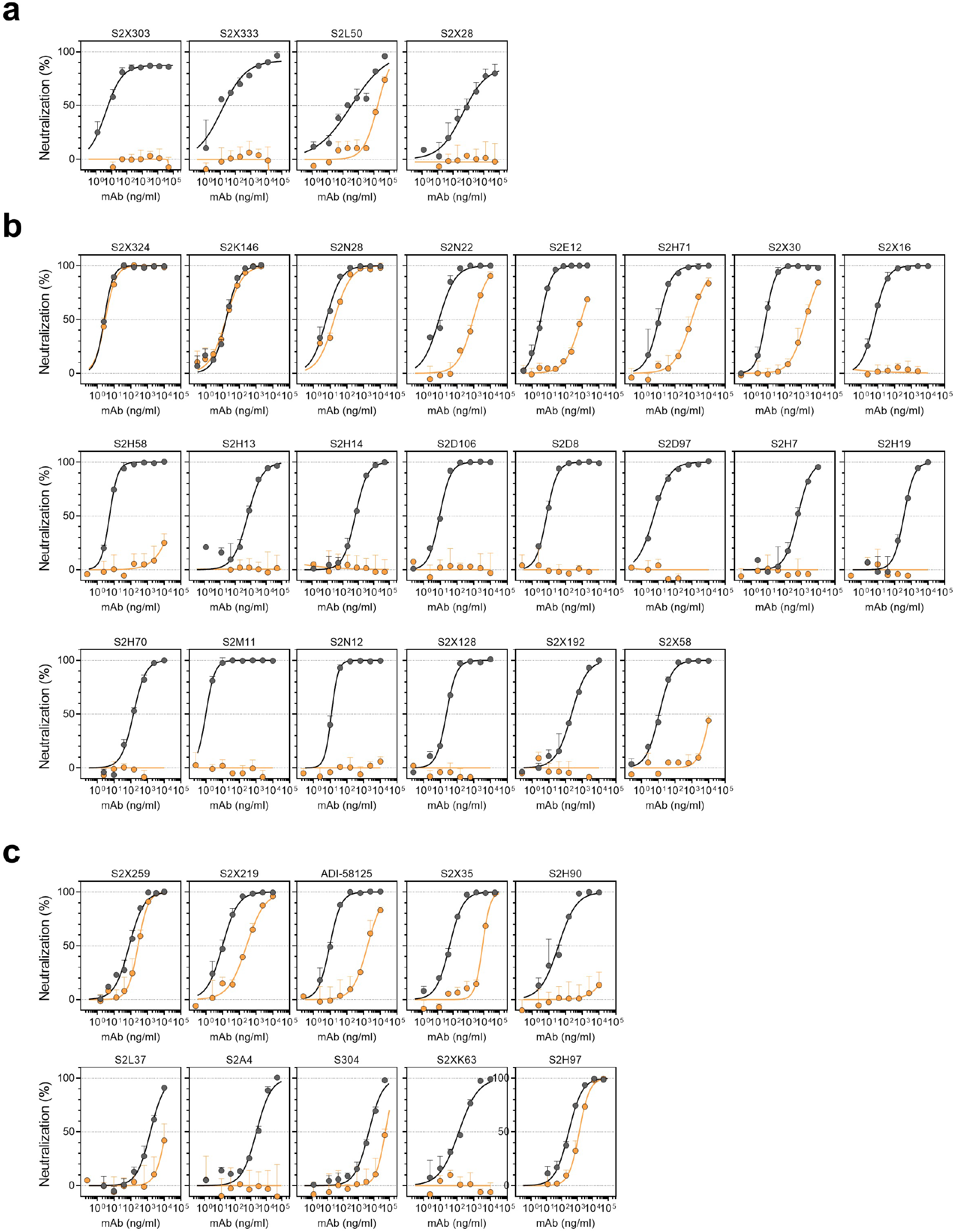
Neutralization of WT (D614) and Omicron SARS-CoV-2 Spike pseudotyped virus by a panel of 36 mAbs. **a-c**, Neutralization of SARS-CoV-2 VSV pseudoviruses carrying wild-type D614 (grey) or Omicron (orange) S protein by NTD-targeting (**a**) and RBD-targeting (**b-c**) mAbs (**b**, site I; **c**, sites II and V). Data are representative of one independent experiment out of two. Shown is the mean ± s.d. of 2 technical replicates.

